# Compost from urban biowaste shapes the bacterial community and its functional traits in the rhizosphere of tomato plants

**DOI:** 10.64898/2026.10.28.762700

**Authors:** Rosalba Cipriani, Nadine De Biasio, Maëva Labouyrie, Giulia Ghirardello, Werner Zanardi, Tiziano Bonato, Barbara Baldan, Sebastiano Nigris

## Abstract

Industrial compost derived from urban biowaste is a sustainable biofertilizer made from food waste and plant matter. It has long been used as a fertilizer in agriculture and horticulture for its rich nutrient content. However, the composition of the microbiota in industrial biowaste compost has been studied only partially, and the effects of this bacterial community on soil, rhizosphere, and plant root communities have received limited attention. In this study, we conducted community and functional analyses to examine how the microbiota in biowaste compost influences bacterial diversity, community composition, and metabolic potential in tomato plants, compared with plants fertilized with conventional, unsustainable chemical fertilizer. Source contribution analysis tracking bacterial taxa indicated a significant contribution of compost-derived taxa to the soil, rhizosphere, and roots. The community analysis revealed distinct clustering by fertilizer type, with biowaste compost promoting a unique, more diverse rhizosphere community, enriched in Firmicutes species. Functional analysis highlighted increased microbial pathways in the rhizosphere treated with biowaste compost, including enzymes for cellulolytic and xylanolytic activity, nitrogenase, phytase, and 1-ACC deaminase. Overall, food-derived industrial biowaste compost enhances bacterial diversity and the predicted functional and plant growth-promoting potential in the rhizosphere of tomato plants, underscoring its potential as a sustainable biofertilizer that supports soil health, microbial biodiversity, and crop productivity.

## 1. INTRODUCTION

Different types of composts have been used for decades as soil amendments and fertilizers, and their benefits for plant growth are well known. These positive effects are largely attributed to its organic matter content, which benefits the soil (Lehmann and Kleber, 2015; Pergola et al., 2018; Zhao et al., 2022), and to its macro- and micronutrient content, particularly nitrogen (Lucchetta et al., 2023). However, little is known about how the microbial community of different compost types interacts with soil and plant-associated microbiotas. These interactions have been investigated, for example, in manure (Guo et al., 2022), in digestates derived from biogas production using livestock and agro-industrial residues (Caradonia et al., 2019), and in composted materials derived from plant-only-based waste (Antoniou et al., 2017). These studies have shown that the use of these soil organic amendments could influence the composition of the soil and plant rhizosphere bacterial communities (Antoniou et al., 2017; Guo et al., 2022). However, these studies are limited to material with specific sources, and do not include industrial biowaste soil amendments. In fact, the environmental benefits of composting are further enhanced when implemented at an industrial scale within a circular-economy framework, thereby integrating composting into urban organic waste management systems. Through a multistep biowaste transformation process, it is possible to generate not only compost but also energy (biogas and heat), carbon dioxide for human consumption, and clean water (Fava et al., 2015). Furthermore, this compost produced from urban organic waste is a carbon-rich material that can be further enriched by incorporating plant residues from agriculture, horticulture, and home gardening. The production and use of biowaste organic fertilizers, such as compost, are attracting growing interest, driven by the rising costs of chemical fertilizers and increasing awareness that microbial biodiversity in agricultural soils is threatened by intensive farming practices, unregulated synthetic fertilizer use, and climate change (Cavicchioli et al., 2019; Jansson and

Hofmockel, 2020; Ibáñez et al., 2023). These environmental stresses can affect the abundance and composition of soil bacterial communities, including those associated with the rhizosphere (the region of the soil in direct contact with roots) and with plant root tissues (rhizodermis and inner tissues), where most beneficial interactions crucial for the plant health occur (Bulgarelli et al., 2013; Philippot et al., 2013; Trivedi et al., 2020; Sokol et al., 2022).

It is known that industrial compost from biowaste compost has its own bacterial microbiota (Tondello et al., 2022; Martínez-Yáñez et al., 2023), however, what is still unclear is if the bacterial species present in the compost, and selected by the industrial process of compost production, may interact with soil and plant bacterial communities. In this study, we hypothesized that biowaste compost produced through an industrial process involving the treatment of municipal food waste and plant residues could: (i) influence the microbial communities associated with soil, the rhizosphere, and the root of tomato plants, in comparison to the effects of conventional chemical fertilization; and (ii) whether differences in the communities of plants grown with compost *versus* plants grown with conventional chemical fertilizers might also be reflectedchemical fertilizers might also be reflected at a functional level, with soil and plant-associated communities under compost fertilization showing a higher metabolic potential.

To test our hypotheses, we employed a well-established 16S rRNA gene metabarcoding approach to characterize and compare bacterial communities in bulk soil, the rhizosphere, and the roots of tomato plants subjected to different fertilization treatments. The corresponding metabolic profiles were inferred from the taxonomic composition of these communities, and functional groups related to plant growth promotion, plant material/organic carbon degradation (i.e. chemoorganotrophic species) and nitrogen usage have been compared. Our analysis revealed that microbial communities clustered distinctly according to the fertilization regime, with industrial biowaste compost application fostering a unique and more diverse rhizosphere bacterial microbiota, and an enrichment of microbial pathways. These findings underscore the potential of industrial biowaste compost as a sustainable biofertilizer, capable of enhancing rhizosphere microbial diversity and metabolic capacity. Taken together, the results of this study, combined with the inherent sustainability of composting food scraps and plant residues, support a data-driven approach to guide sustainable management practices in both urban waste recycling and agricultural soil amendment. This work highlights the potential of compost as an ecologically sound alternative to conventional inputs, contributing to integrated strategies for circular economy and soil health preservation.

## 2. MATERIALS AND METHODS

*Abbreviations used in the text:*

Samples analysed: BS=bulk soil; C=only-compost; RH=rhizosphere; R=root; SOIL=unamended-soil.

Treatments analysed: M42=compost; M42ST=sterilized compost; U=urea.

### 2.1 Soil and compost used for plant cultivation

The soil used for the experiment consisted of a mixture of 45% agricultural field soil, 45% peat, 5% perlite, and 5% sand. Its chemical and physical properties have been determined according to the UNI standard methods (Table S1). The compost used in this work was produced by S.E.S.A. S.p.A. (Este, Italy), and the physical-chemical characteristics were determined according to UNI standard methods and were in line with EU directives and consistent with the company standards (Table S2). The sterile compost was obtained by autoclaving at 121°C for 2 h, and its sterility was verified as follows: 10 g of sterile compost were added to 50 mL of sterilized deionized water and left under agitation on the orbital shaker for 1 hour. 100 µL of the solution was plated on PCA agar with the spread plate method and incubated at 28°C for 4 days. After this period, no colonies were observed on the plate, confirming successful sterilization. To verify any possible effect of the sterilization process on plant substrate, the pH of soil/compost and soil/sterile compost mixtures was measured, with no significant difference observed.

### 2.2 Experimental design for community analysis

Twenty plants of *Solanum lycopersicum* L. cv. Micro-Tom were grown under controlled conditions in pots filled with soil supplemented with three treatments: 33 g/L of compost, 33 g/L of sterile compost, or 0.36 g/L of urea. The amount of compost to be used in the experiment was calculated based on the nitrogen (N) requirement of tomato plants, which is 150 kg N/ha. Urea has been chosen as a chemical fertilizer due to its nitrogen release rate being comparable to that of compost (Qu et al., 2020).

### 2.3 Plant cultivation and growth

Tomato seeds (*Solanum lycopersicum* L.) cv. Micro-Tom were surface sterilized using 5% sodium hypochlorite on the orbital shaker for 20 minutes, followed by 4 washings with sterile water. After 48 hours of imbibition at RT, seeds were sown on half-strength Murashige & Skoog Basic Medium supplemented with 1.5% agar, and germinated at 25 °C, 65% humidity, and a photoperiod of 16 h light. Five days after germination, seedlings were transplanted into 0.5 L pots filled with soil supplemented with the respective treatment and placed back in the growth chamber under the same conditions, with a random disposition to avoid position-dependent bias. All the plants and pots were watered evenly and regularly for 5 weeks to allow biomass development.

### 2.4 Sampling for metabarcoding analysis and for biomass measurements

Bulk soil (BS), rhizosphere (RH), roots (R) and leaves from 10 randomly chosen plants for each treatment were collected aseptically to avoid cross-contamination. Bulk soil samples were collected as far as possible from the roots. To collect rhizosphere, roots were shaken to remove loosely attached soil particles, then vortexed in a 50 mL tube to detach and collect rhizosphere soil particles. Roots were then washed to remove any rhizosphere soil particles, gently dried with sterile paper, and immediately frozen. Samples from Biowaste compost and soil were also taken to characterize their bacterial community.

Leaves from plants treated with compost, sterile compost and urea were dried for five days at 60 °C and weighed on an analytical balance to assess their dry biomass. Statistically significant differences in dry biomass between treatments were tested with the Kruskal-Wallis rank sum test and groups were then compared in pairs with a pairwise Wilcoxon rank-sum test. Resulting p-values were adjusted with the Benjamini-Hochberg (BH) method.

### 2.5 DNA extraction and sequencing

DNA was extracted from bulk soil, rhizosphere, and roots treated with biowaste compost M42 (BS.M42, RH.M42, R.M42), sterile biowaste compost M42 (BS.M42ST, RH.M42ST, R.M42ST), and urea (BS.U, RH.U, R.U), unamended soil (S), and from compost M42 (C.M42). DNeasy Power Soil Kit (Qiagen) was used for DNA extraction, following the manufacturer’s protocol.

DNA concentration and purity were then quantified by a Nanodrop spectrophotometer (NanoPhotometer®, Implen). The target region V5-V7 of the 16S rRNA gene was amplified using primers 799F (5’AACMGGATTAGATACCCKG3’) and 1193R (5’ACGTCATCCCCACCTTCC3’), to minimize contamination from plant sequences. Amplicons were then used for amplicon library preparation, which involved end-repair, A-tailing, and ligation with Illumina adapters, followed by library quality control. Samples were sequenced on an Illumina NovaSeq 6000 platform by Novogene Europe (UK), producing PE 250 bp raw reads.

### 2.6 Community analysis

Demultiplexed paired-end fastq files were processed in R v.4.3.0, using the DADA2 R script (Callahan et al., 2016). The error rates were used by the DADA2 algorithm to infer the true sequence variants from the initial unique sequences. Forward and reverse reads were merged to obtain the full denoised sequences, and an Amplicon Sequence Variant (ASV) table was created. After chimeric ASV identification, taxonomy was assigned to each ASV. The absolute abundance of ASVs was normalized using a standard of sequence number corresponding to the sample with the fewest sequences. Subsequent analyses of alpha diversity and beta diversity were performed based on normalized data. A rarefaction curve was obtained to assess whether the sequencing depth was sufficient to describe the diversity of the communities. This was made by subsampling the sequences at increasing depths and recording the number of observed ASV. Relative abundance, alpha, and beta diversity analyses were performed using the R package *phyloseq* v1.44.0 (McMurdie and Holmes, 2013). Statistical analyses (ADONIS, Kruskal-Wallis, T-test) were performed using the R *vegan* package. The functional potential of the microbial communities was investigated using the R package *microeco* based on the FAPROTAX database (Liu et al., 2021). Statistical significance was assessed by ANOVA analysis and Tukey test. For this analysis, functional categories associated with organic matter, plant material, and nitrogen transformation were selected, according to Labouyrie et al. (2023). Metabolic pathways and enzymatic activities were predicted using PICRUSt2 ((Douglas et al., 2020)). Samples from the same origin were analyzed together as biological replicates, and results are presented as groups of samples (BS, RH, R, S, C).

### 2.7 Bacterial taxa dynamics analysis

The potential transfer of bacterial taxa from compost and soil to plant-associated environments was investigated using FEAST in the R environment (Shenhav et al., 2019). This software was used to estimate the proportions of bacterial taxa in different microbial communities that were putatively derived from compost or soil sources. All samples (bulk soil, rhizosphere, and roots) were treated as sinks, while compost and soil were considered sources. The analysis was conducted on plants grown with compost and sterile compost. FEAST was run using all ASVs without aggregation at any taxonomic rank, starting from the final ASV table generated with the *phyloseq* package. Statistical analyses of the resulting contribution matrix were also performed in R. Data normality was assessed using the Shapiro–Wilk test applied to the residuals of a linear model, and homogeneity of variances was evaluated using Levene’s test. Because the data did not meet the assumption of normality, group differences were tested using the Wilcoxon rank-sum test. In boxplots, significant pairwise differences (adjusted p < 0.05) were indicated using asterisks.

### 2.8 Statistical analysis of associations between leaf biomass and bacterial families

Associations between leaf dry biomass and bacterial family relative abundance were evaluated using Spearman’s rank correlation. Analyses were performed separately for bulk soil, rhizosphere, and root samples and for each treatment. The 30 most abundant families were selected independently within each compartment. P-values were corrected using the Benjamini-Hochberg method within each compartment and treatment, with adjusted p-values < 0.05 considered significant.

## 3. RESULTS

### 3.1 Different soil fertilization leads to different bacterial communities in the rhizosphere

To understand the differences between the communities, we calculated a matrix of dissimilarities (Non-metric Multidimensional Scaling analysis - NMDS) using the Bray-Curtis (Fig. 1A) and Jaccard (Fig. 1B) to account for differences in species abundance (Bray-Curtis) and in species presence/absence (Jaccard). The NMDS analysis based on Bray-Curtis dissimilarities reveals a clear clustering of samples, reflecting distinct community compositions among the groups. As expected, biowaste compost (C.M42) formed a well-defined and isolated cluster.

**Figure 1.**
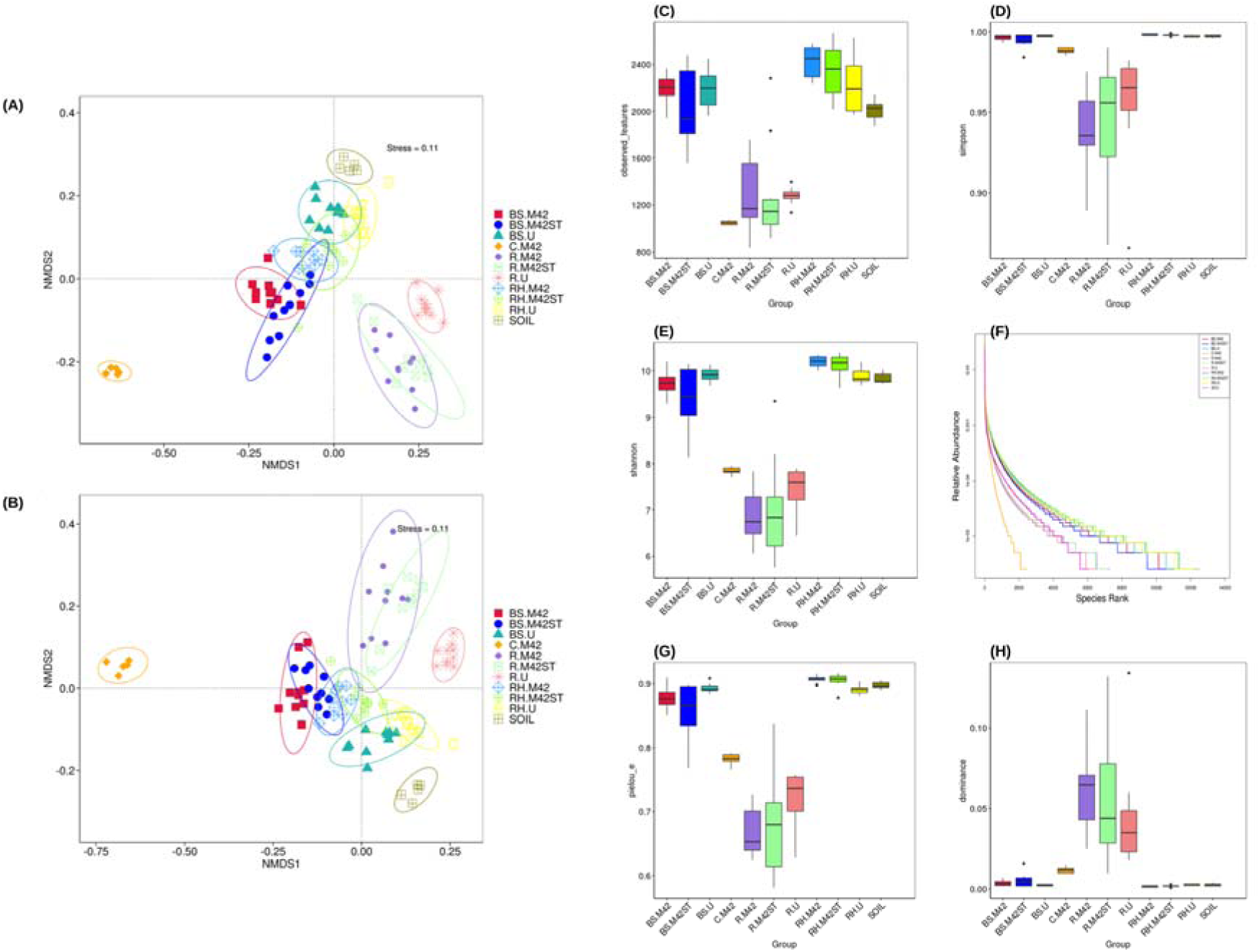
Bacterial community structure, diversity and evenness across sample groups. (A, B) Non-metric multidimensional scaling (NMDS) ordination of bacterial communities based on (A) Bray-Curtis and (B) Jaccard distances (stress = 0.11 for both). Each point is one sample, and ellipses enclose the samples of each group. (C–E) Alpha diversity indices: (C) observed features, (D) Simpson index and (E) Shannon index. (F) Rank-abundance curves showing the relative abundance of ASVs ranked from most to least abundant in each sample group. (G) Pielou evenness index. (H) Dominance index.

The rhizosphere samples (RH.M42, RH.M42ST, and RH.U), exhibited notable clustering patterns with some overlap. The rhizosphere of plants fertilized with sterilized biowaste compost (RH.M42ST) and the rhizosphere of plants fertilized with urea (RH.U) are close to each other, indicating compositional similarity. In contrast, the rhizosphere of plants fertilized with biowaste compost (RH.M42) formed a distinct cluster with minimal overlap, indicating a more distinct community composition than the other RH groups.

The R² values from the ADONIS analysis (Table S3) support these findings: the rhizosphere bacterial community of plants fertilized with sterilized biowaste compost was more similar to that of urea-fertilized plants (R² RH.M42ST–RH.U = 0.10) than was the community of plants fertilized with untreated biowaste compost (R² RH.M42–RH.U = 0.13). Root communities (R.M42, R.M42ST and R.U) clustered tightly and were clearly separated from rhizosphere communities (R² = 0.13–0.19) and bulk soil communities (R² = 0.15–0.22).

The roots of plants fertilized with biowaste compost (R.M42) exhibited relatively low within-group variability. The roots of plants fertilized with sterilized biowaste compost (R.M42ST) showed slight overlap with roots treated with urea (R.U), suggesting some shared species. Among the various root sample groups, the R² values from the ADONIS analysis are generally lower, but the trend observed in rhizosphere samples is also detectable in root samples (Table S3).

Samples from bulk soil groups (BS.M42, BS.M42ST, and BS.U) exhibited some variability in clustering patterns. The bulk soil treated with biowaste compost (BS.M42) group displayed a compact cluster with minimal overlap, indicating a relatively homogeneous community composition. Interestingly, the bulk soil added with sterilized biowaste compost (BS.M42ST) group showed moderate overlap with bulk soil treated with urea (BS.U, Fig 1A), suggesting that these two communities might be more influenced by the soil community, and,In contrast, that bacterial species from biowaste compost may have contributed to shaping the BS.M42 community.

Similar results have been obtained by conducting ADONIS analysis on Jaccard distance matrix (Table S4), confirming the community differences highlighted by the previous analysis. These results are further supported by the PCoA visualizations of Bray-Curtis and Jaccard analyses (FigS1A and B), as well as by NMDS and PCoA that consider the compartments separately (FigS1C and D).

Rarefaction analysis (FigS2) revealed that the sequencing depth was adequate to capture community diversity. To investigate the communities in terms of species richness and diversity, we calculated three indexes: observed features, the Simpson index, and the Shannon index (Fig. 1 C, D, E).

All the indexes indicate that the rhizosphere and bulk soil samples have the highest number of taxa, compared with root samples, which have fewer taxa, as expected in an environment under stringent plant control. Interestingly, biowaste compost samples (C.M42) exhibited alpha diversity indexes lower than those of the soil, with a community that is less evenly distributed than the soil community. Comparing the statistical significance of the various group comparisons for the considered indexes (TableS5), we found that the rhizosphere of plants fertilized with biowaste compost (RH.M42) has a higher alpha diversity than the rhizosphere of plants fertilized with urea (RH.U), despite an uneven distribution of species. Interestingly, the rhizosphere of plants fertilized with sterile biowaste compost presents higher indexes for alpha diversity compared to RH.U and lower indexes compared to RH.M42, showing statistical significance only for the Simpson and Shannon indexes (RH.M42ST vs RH.U) and no significance when compared to RH.M42.

No significant differences were detected within the bulk soil group, and, surprisingly, no bulk soil samples differed significantly from soil samples. Notably, BS.U samples show no significant difference from RH.U, yet this result is challenging to interpret, whereas both BS.M42 and BS.M42ST differ from their respective rhizosphere samples. As with bulk soil samples, root samples showed no significant differences across fertilization types.

To describe the distribution of species in the samples, we calculated the relative abundance, Pielou index, and dominance index. The relative abundance plot (Fig. 1F) shows that C.M42 compost exhibits the most uneven community, with fewer species unevenly distributed. In contrast, samples from the rhizosphere of plants fertilized with biowaste compost and from plants treated with sterile biowaste compost have a higher number of rarer, more evenly distributed taxa.

The Pielou index (Fig. 1G) indicates that bulk soil and rhizosphere samples host equally distributed communities in terms of abundance. Bulk soil with urea shows a significantly more even community than bulk soil with biowaste compost or sterile biowaste compost. Interestingly, the rhizosphere with compost and the rhizosphere with sterile compost exhibit significantly higher Pielou indices, indicating more even communities than the rhizosphere with urea. In contrast, plant roots are colonized by unevenly distributed communities compared with the bulk soil and rhizosphere compartments, and this pattern is more pronounced in roots with biowaste compost and sterile biowaste compost. Although not statistically significant, the Dominance analysis (Fig. 1H) further supports this finding, showing that there are few dominant species in the roots compared with the other communities analyzed in this work. Consistent with the Pielou index analysis, RH.M42 and RH.M42ST have significantly lower Dominance indices, indicating fewer dominant species than RH.U. The statistical significance of the comparisons among groups for the Pielou and Dominance indices is reported in TableS6.

### 3.2 Taxa distribution and relative abundance differ based on the fertilization treatment

The analysis of the relative abundance of bacterial phyla (Table S7), showed that within the ten most representative phyla, Proteobacteria, Actinobacteriota, Bacteroidota, and Firmicutes are the most dominant (Fig. 2). Biowaste compost shows a relatively simple community dominated by species from the phylum Firmicutes. Proteobacteria are also present, albeit to a lesser extent than in the soil. Family-level relative abundance (Fig S3) indicates that the most prevalent family in biowaste compost is Bacillaceae (Table S8).

**Figure 2:**
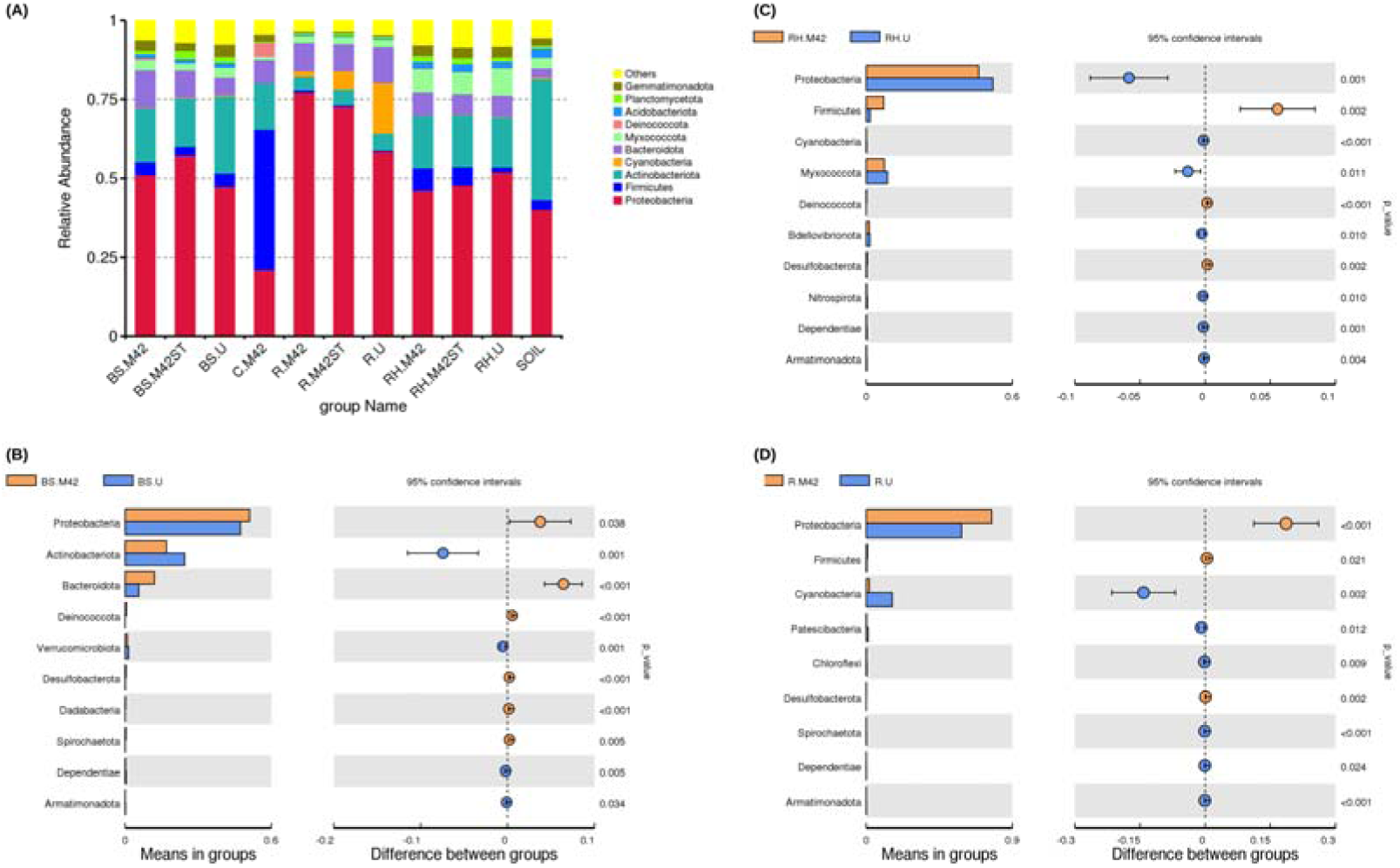
Description and statistical analysis of relative abundance and phyla distribution among sample groups. In **(A)** is the barplot of relative abundance and distribution of the first 10 phyla; **(B)** Phyla enrichment in compost vs urea samples in bulk soil, **(C)** rhizosphere, and **(D)** roots. Bars represent mean values of phylum abundance in the group. Statistical significance was detected by Student’s *t*-test, and the relative P values are reported in the plot axis.

If we compare the phylum composition of the rhizosphere with biowaste compost and with urea (Fig. 2A), we observe that the ratio of enriched phyla differs from that of the bulk soil. RH.M42 exhibited only three enriched phyla when biowaste compost is used (Fig. 2C), whereas seven phyla were identified when urea is applied as fertilizer. Notably, in RH.M42 a strong enrichment is evident for species belonging to the Firmicutes phylum, confirming the distribution observed in all the plants fertilized with biowaste compost (Fig. 2A). Families Bacillaceae and Halomonadaceae are enriched in the biowaste compost-fertilized samples (Fig. S4B, Table S10). In contrast, we observed an enrichment of species belonging to Nitrosomonadaceae, and Nitrospiraceae families in urea-treated samples.

Comparing the phylum composition of bulk soil with biowaste compost and with urea (Fig. 2B) reveals that bulk soil with biowaste compost is enriched in six phyla compared to the one with urea: Proteobacteria, Bacteroidota, Deinococcota, Desulfobacterota, Dadabacteria, and Spirochaetota. Bulk soil fertilized with biowaste compost has five phyla significantly enriched compared to the bulk soil fertilized using sterile biowaste compost (Fig. 2A), Bacteroidota, Myxococcota, Deinococcota, Gemmatimonadota, and Dadabacteria.

Many bacterial families (47 out of 93) are enriched in the bulk soil with biowaste compost compared to the bulk soil with urea (Fig. S4A and Table S9), indicating a community shift when biowaste compost is utilized.

Phylum-level enrichment analysis of root samples from plants fertilized with biowaste compost and with urea (R.M42 vs R.U) revealed that, consistent with the bulk soil and the rhizosphere, R.M42 was enriched in Firmicutes and Proteobacteria (Fig. 2D). At the family level, biowaste compost-treated roots showed increased abundance of Rhizobiaceae, Pseudomonadaceae, Comamonadaceae, and Bacillaceae (Fig S5A and Table S11).

### 3.3 Biowaste compost fertilization leads to a higher presence of Firmicutes genera in all the samples

Composition and relative abundance analyses indicate that the bacterial community in biowaste compost is particularly enriched in Firmicutes. By examining the distribution of Firmicutes genera across various sample groups and the distribution and relative abundance of the top 20 Firmicutes genera across all samples (Fig. S6), we detected enrichment of 5 Firmicutes genera in the rhizosphere of plants fertilized with biowaste compost compared with samples from plants fertilized with urea (Fig. S5B and Table S10). This number drops to zero when comparing the rhizosphere of plants fertilized with sterile biowaste compost with that of plants fertilized with urea (Fig. S7 and Table S12).

### 3.4 Bacterial source analysis supports a direct transfer of bacterial taxa from compost to rhizosphere, roots, and bulk soil

To test our hypothesis of a direct transfer of bacterial taxa from compost to the bulk soil, rhizosphere, and plant roots, we used FEAST software comparing samples from the three compartments (BS, RH, and R) fertilized with biowaste compost to those fertilized with the same compost that was sterilized.

The source tracking analysis (Table S13; Fig. 3) showed that the proportion of bacterial taxa originating from compost was significantly higher in the rhizosphere (RH.M42), roots (R.M42), and bulk soil (BS.M42) compared with the corresponding samples fertilized with sterile compost, in which the microbial component had been eliminated by sterilization. Notably, the rhizosphere hosted the highest proportion of compost-derived bacterial taxa, reinforcing its role as a key reservoir of microorganisms with the potential to interact with plant roots.

**Figure 3:**
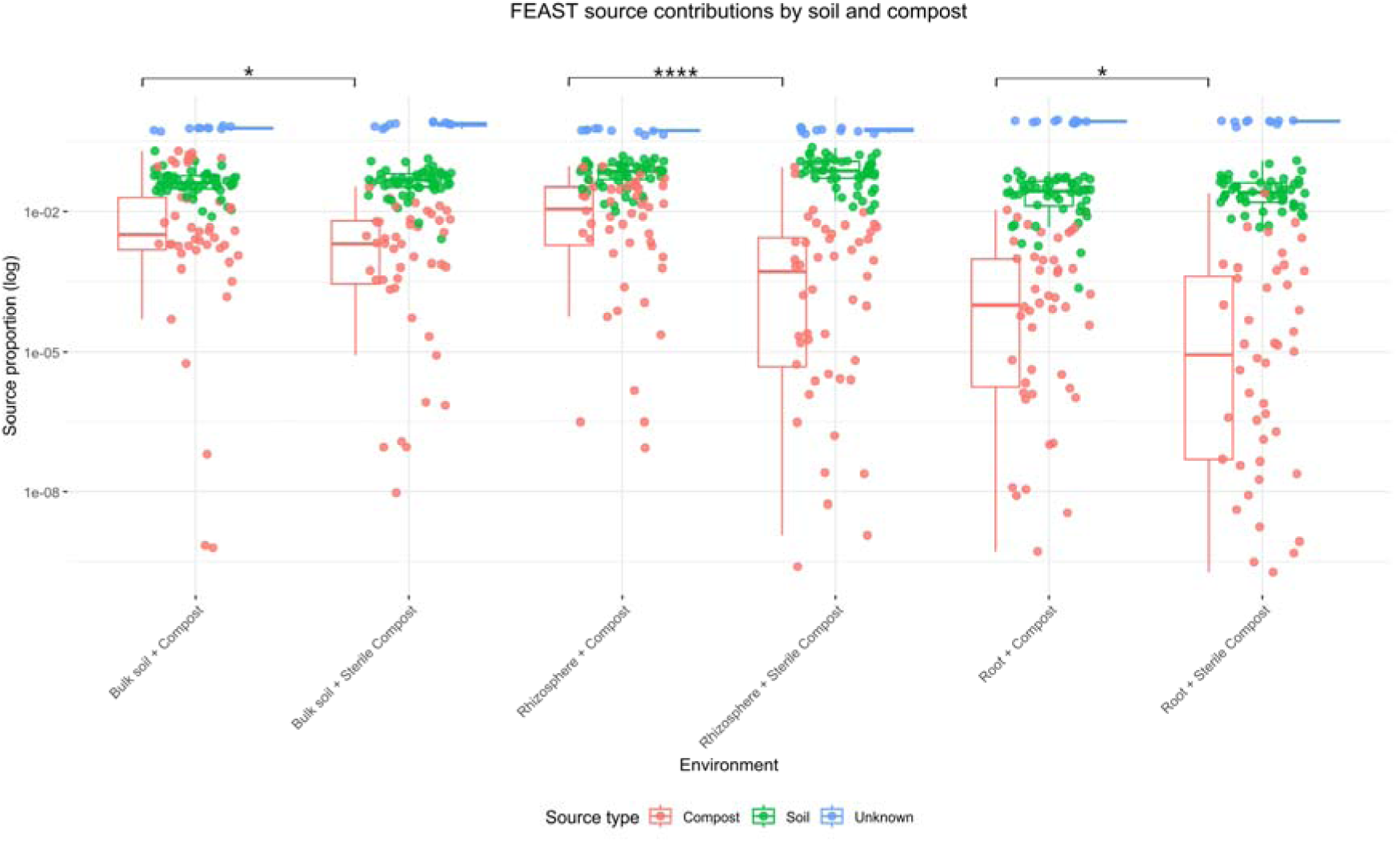
FEAST analysis of source contributions to the group of samples (sinks). Boxplots describe the contribution (source proportion, log values) of the sources (Compost C.M42 and SOIL) to the community of each sink (Bulk Soil, Rhizosphere, Roots) from plants cultivated with Compost and sterilized compost. Group statistical differences were tested using the Wilcoxon rank-sum test. In boxplots, significant pairwise differences were indicated using asterisks (* adj P < 0.05; **** adj P < 0.0001).

### 3.5 The use of biowaste compost results in variations in microbial functional traits

To assess whether applying biowaste compost could alter functional traits in the samples, we conducted a functional analysis of community composition. In this context, we measured the relative abundance of species categorized as chemoheterotrophs, nitrogen fixers, nitrifying bacteria, denitrifying bacteria, cellulolytic bacteria, chitinolytic bacteria, xylanolytic bacteria, and human-associated bacteria (Fig. 4). The results indicated that bulk soil with biowaste compost had a higher proportion of xylanolytic bacteria than bulk soil with urea (Fig. 4G). In contrast, urea application increased the proportions of nitrogen fixers, nitrifying bacteria, and chitinolytic bacteria compared with both compost treatments (Fig. 4B, C, and F). Denitrifying bacteria showed the opposite pattern, being significantly more abundant in bulk soil amended with sterile biowaste compost than in urea-treated soil (Fig. 4D). In the rhizosphere, biowaste compost application increased the proportion of cellulolytic bacteria (Fig. 4E) and xylanolytic bacteria (Fig. 4G). In contrast, urea application resulted in higher proportions of chemoheterotrophic, nitrifying, denitrifying, and chitinolytic bacteria (Fig. 4A, C, D, and F). In root samples, we observed a trend toward higher chemoheterotrophs, supported by a significant rise when sterile biowaste compost was added to the soil (Fig. 4A). Conversely, urea application resulted in statistically significant increases in the percentages of nitrogen fixers, denitrifying bacteria, cellulolytic bacteria, and chitinolytic bacteria (Fig. 4B, D, E, and F), whereas nitrifying bacteria were barely detectable in roots and did not differ among treatments (Fig. 4C). These data also suggest that none of the fertilization regimes we employed in this test significantly increased the presence of human-associated bacteria, despite biowaste compost being a fertilizer derived from human domestic biowaste (Fig. 4H)

**Figure 4:**
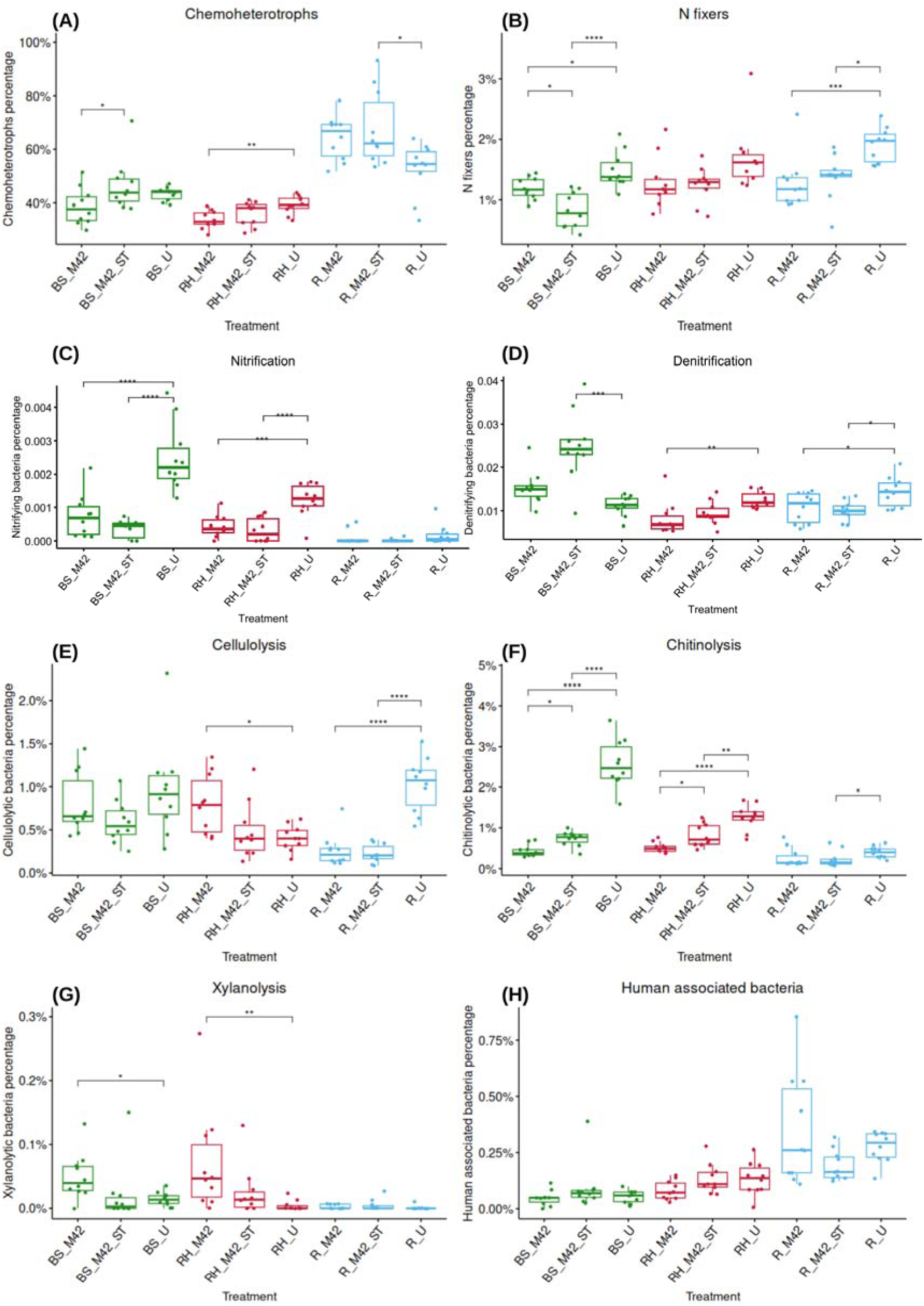
Functional analysis of microbial traits. Comparison among the diverse treatments applied for evaluating the relative abundance of species categorized as chemoheterotrophs **(A)**, nitrogen fixers **(B)**, nitrifying bacteria **(C),** denitrifying bacteria **(D)**, cellulolytic bacteria **(E)**, chitinolytic bacteria **(F)**, xylanolytic bacteria **(G)**, and human-associated bacteria **(H)**. Values represent means ± SD. Statistical significance was assessed by ANOVA analysis and Tukey test. *P < 0.05, ** P < 0.01, *** P < 0.001

From the community composition, it is also possible to infer potential metabolic pathways and how they change across the different fertilization modalities (Fig. 5). Enriched pathways in biowaste compost-fertilized samples, compared with urea-fertilized samples, are shown as red dots in the volcano plots. In bulk soil and roots, we did not observe a significant increase in these pathways in biowaste compost-fertilized samples (Fig. 5A and 5C, Table S14 and Table S15). Conversely, in the rhizosphere of biowaste compost-enriched samples (Fig. 5B, Table S16), we observed a marked increase in these pathways, indicating that this environment may be more metabolically active in plants fertilized with biowaste compost. Pathway annotations are provided in Table S17. These potential metabolic activities have been further investigated by analyzing the potential enzymatic activities associated with the different community species (Table S18). Specific enzymes linked to plant-promoting activities have been examined in the rhizosphere, revealing that biowaste compost samples are significantly enriched in species with potential enzymatic activities: dihydroxyacetone phosphate aldolase (involved in carbon fixation and sugar metabolism), nitrite reductase (ammonia-forming, reduces nitrite to ammonia), phosphoribosyltransferase (involved in organic phosphate metabolism), lipase (degrades pathogenic biofilms and lipid structures), phytase (releases inorganic phosphate from phytates), nitrogenase (fixes atmospheric nitrogen into ammonia), gibberellin 3-beta-dioxygenase (synthesizes gibberellins), - glutamate synthase (important in nitrogen assimilation), acid phosphatase (hydrolyzes organic phosphorus compounds), 1-Aminocyclopropane-1-carboxylate deaminase (lowers ethylene levels in plants), siderophore biosynthesis, and polymer degrading enzymes such as cellulase, xylanase and pectate lyase. EC codes of potential PGP enzymatic activities are reported in Table S18.

**Figure 5:**
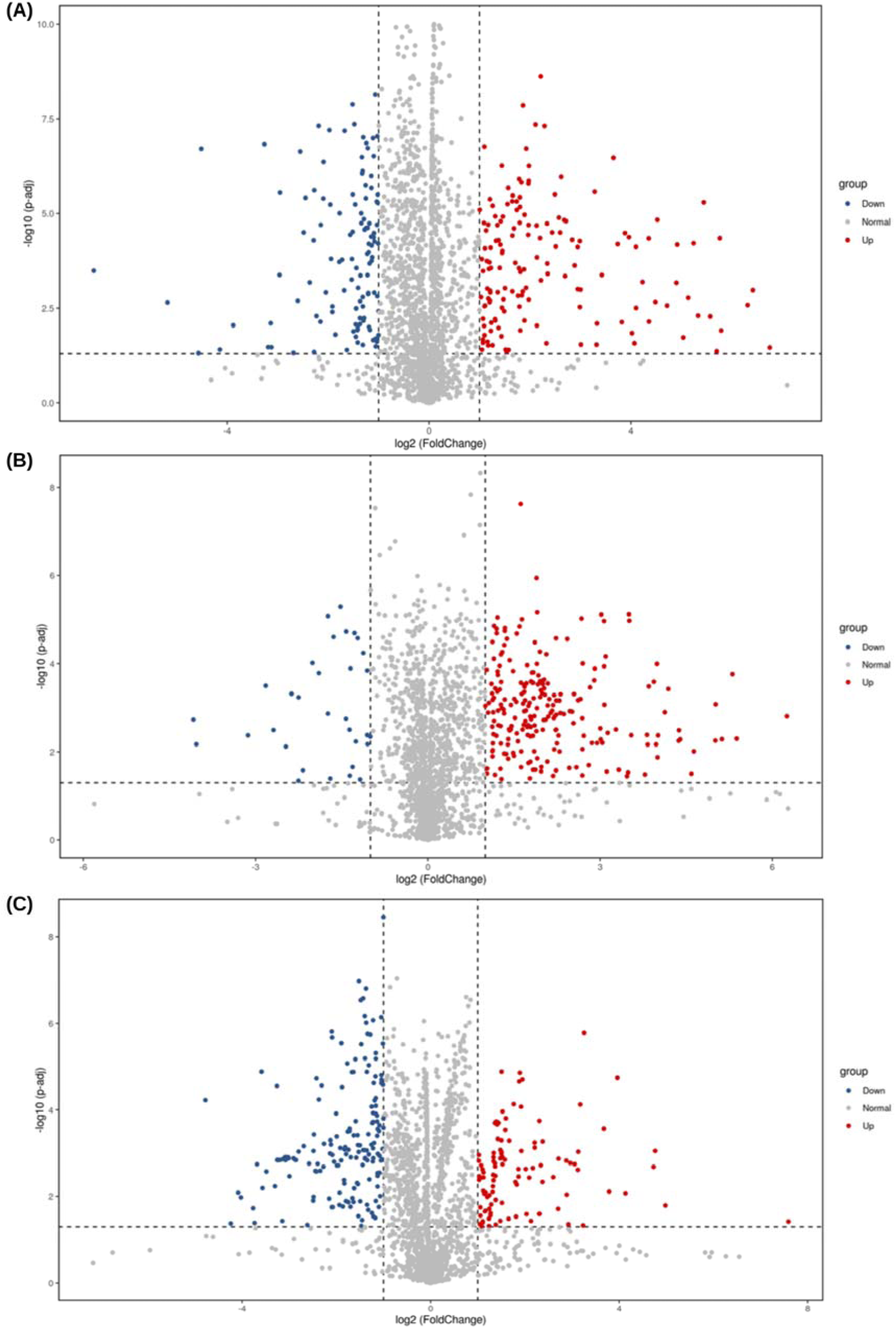
Enrichment analysis of potential metabolic pathways. Volcano plot of the pathway enrichment analysis between sample groups: **(A)** BS.M42 vs BS.U; **(B)** RH.M42 vs RH.U; and **(C)** R.M42 vs R.U. Plots show only data significantly different. Each point in the figure represents a function with differences, where ‘Up’ represents that the abundance of the function with differences in the first comparison group is higher than that in the second, while ‘Down’ represents the opposite. Statistical significance was assessed by Student’s t-test, on the Y-axis is the P-value of the significance test

### 3.6 Associations between bacterial family abundance and tomato leaf biomass

To investigate whether the compost-associated bacterial community was related to the physiological response of tomato plants, leaf dry biomass was measured in the plants used for the community experiment and correlated with the relative abundance of bacterial families in the rhizosphere and root compartments (Table S19).

Leaf biomass measurements showed that plants treated with compost or sterile compost maintained significantly higher biomass than those treated with urea (Fig. S8A).

We subsequently assessed the correlations between leaf biomass and the relative abundance of bacterial families. In the rhizosphere, it was possible to observe some trends, although they were not statistically significant. Under compost treatment, the strongest positive associations, based on the lowest p-values, were observed for TRA3-20, Saprospiraceae, and Reyranellaceae. Under sterile compost, the lowest p-values for positive associations were observed for TRA3-20, Flavobacteriaceae, and Reyranellaceae. Under urea treatment, Pedosphaeraceae, Reyranellaceae, and Devosiaceae showed the lowest p-values among the positive associations (Fig. S8B).

In the root compartment, the three positive associations with the lowest p-values under the compost treatment involved Rhizobiaceae, Microbacteriaceae, and Caulobacteraceae. Under sterile compost, Methylophilaceae, Flavobacteriaceae, and Pseudomonadaceae showed the lowest p-values, with Methylophilaceae being the only statistically significant positive association. Microbacteriaceae was the only family showing a significant negative association under sterile compost. Under urea treatment, the lowest p-values were observed for Saprospiraceae, Microbacteriaceae, and Microscillaceae, although none of these associations was statistically significant (Fig. S8C).

## 4. DISCUSSION

Industrial biowaste compost, produced from the biological treatment of organic waste, mostly food waste and gardening materials, has been used for decades because of its nutrient content (i.e., carbon and nitrogen). However, unlike manure and manure-derived biowaste composts, the microbiota of industrial biowaste compost and its effects on soil and plant-associated communities have been only partially studied, and never compared with the use of chemical fertilizers. To fill this gap, this work aimed to assess the effects of this sustainable fertilization on soil and plant-associated bacterial communities that may benefit the plant, rather than evaluate the impact of biowaste compost on plant growth and yield.

The results of this study highlight the significant impact of biowaste compost application on the composition, diversity, and functional potential of bacterial communities in the rhizosphere, bulk soil, and roots of tomato plants, compared with plants grown with chemical fertilizer (Qu et al., 2020). Our findings reveal the establishment of distinct rhizosphere communities that vary by fertilization treatment.

The analysis of taxonomic groups showed a positive trend in bacterial diversity associated with biowaste compost application in the rhizosphere, an effect that can be leveraged to counteract the deleterious impact of climate change on soil and rhizosphere microbial biodiversity. Notably, the rhizosphere of biowaste compost-treated plants showed the highest median richness and diversity, and the community dynamics analysis indicated that biowaste compost introduces new bacterial species. Biowaste compost application might also stimulate the proliferation of dormant or previously undetectable species: this effect may be attributed to the introduction of organic matter, which serves as a nutrient source for bacteria. The fertilization effect on bacterial species has also been observed in previous work, where the application of biowaste compost to grapevines cultivated in vitro stimulated the root microbiota, increasing the proliferation of rare taxa (Tondello et al., 2022).

The alpha diversity analysis revealed a progressive decrease in the number and evenness of taxa from the bulk soil to the root, highlighting the increasing selective pressures exerted by plants on microbial colonization (Bulgarelli et al., 2013; Xiong et al., 2021). However, the biowaste compost-treated rhizosphere consistently exhibited higher alpha diversity and more even bacterial communities than the urea-treated rhizosphere. This enrichment is likely due to the direct delivery of taxa and the influence of biowaste compost on microbial recruitment, stimulating bacterial taxa that can thrive in the rhizosphere. The dominance of Firmicutes in biowaste compost and biowaste compost-treated plant samples suggests a possible transfer of species from the biowaste compost to plant-associated tissue, particularly the rhizosphere. In this nutrient-rich, complex environment, bacterial species derived from biowaste compost may find favourable conditions for thriving. This hypothesis is supported by results from plants fertilized with previously sterilized biowaste compost: eliminating the living compost microbiota led to a less clear delivery effect in the analyzed samples. Moreover, in samples fertilized with biowaste compost, we observed partial enrichment of species belonging to Bacillaceae, Rhizobiaceae, and Halomonadaceae, families that play critical roles in nitrogen fixation, biocontrol, and plant stress resilience, with the latter family linked to resistance to saline stress (Li et al., 2020). Interestingly, the family Comamonadaceae, enriched in the roots of plants fertilized with biowaste compost, includes several genera, among which the genus Variovorax is a key player in the tomato root microbiota. It has been demonstrated that species belonging to the genus Variovorax are essential for modulating and mediating the effects of other species on root development, protecting the plant from extreme or undesirable effects, and maintaining a high level of complexity and, consequently, high biodiversity within the tomato root community (Finkel et al., 2020; Shi et al., 2023). The source of the enrichment of taxa and the changes in abundance in the rhizosphere and other compartments have been tracked by source enrichment analysis using FEAST. This analysis, which compared samples containing compost with those containing sterile compost, indicates a significant contribution of compost-derived taxa to the samples (bulk soil, rhizosphere, and roots). Evidence of the transfer of taxa from compost to the rhizosphere has been suggested (Guarino et al., 2020; Bellini et al., 2020; Castellano-Hinojosa et al., 2023; Wang et al., 2026), but without computational support from a source composition analysis. Moreover, our analysis also supports a significant transfer of taxa in the root environment, suggesting a higher impact of compost fertilization on the entire microbial community of the plant. This aspect supports our perspective of considering biowaste compost a bio-fertilizer, as it introduces new taxa into the soil and to the plant, thereby enabling the plant to select more bacterial species to interact with.

The differences in the composition of soil and plant-associated bacterial communities may also be reflected in the functional traits and ecosystem services that the bacterial species provide to the environment (Wagg et al., 2014).

The functional analysis of bacterial communities revealed distinct patterns in metabolic potential across treatments. In the rhizosphere, biowaste compost application enhanced the abundance of bacterial species specialized in degrading plant material, such as cellulose and other complex polysaccharides. This may be explained by considering the production process of biowaste compost, which includes a primary anaerobic fermentation step of organic urban biowaste to produce biogas, followed by the addition of plant-driving waste for aerobic digestion to produce biowaste compost. These results, based on inferring ecosystem functions from microbial community composition, present some limitations, as discussed by Labouyrie et al. (2023), but pave the way to investigate these functional/metabolic aspects with advanced omics techniques, such as metatranscriptomics (Romanowicz et al., 2016) and metaproteomics (Fernandes et al., 2022).

The enrichment of potential plant growth-promoting enzymes in biowaste compost-treated rhizospheres further highlights the functional advantages of biowaste compost application. Enzymes such as nitrogenase, gibberellin 3-beta-dioxygenase, and 1-aminocyclopropane-1-carboxylate (ACC) deaminase were significantly enriched in the rhizosphere samples, indicating a potential beneficial activity of the microbial community to support plant growth through nutrient acquisition, hormone production, and stress mitigation by the activity of ACC deaminase, extending as a second genome the host metabolic potential and resilience in stress (Henry et al., 2021). Overall, despite the known limitations of a computational approach, this analysis may also represent a cost-effective strategy for identifying potential metabolic processes associated with fertilization practices and field management strategies for a microbe-friendly agricultural systems (Hartmann and Six, 2023).

Finally, the relationships between leaf biomass and bacterial relative abundance differed according to both plant compartment and treatment. In the rhizosphere, the recurrent positive trend involving Reyranellaceae across all three treatments may suggest an association with microbial processes supporting plant health. Members of this family have previously been linked to soil organic carbon and nitrogen dynamics following compost and biochar application (Bello et al., 2023). A similar role in nutrient cycling has been predicted for

TRA3-20 (Huang et al., 2024), which was positively associated with biomass under both compost and sterile compost treatments. Clearer treatment-specific associations were observed in the root compartment. In the compost treatment, Rhizobiaceae showed the strongest positive, although non-significant, association with leaf biomass, potentially reflecting the recruitment of beneficial root-associated bacteria by more productive plants. Interestingly, under sterile compost, Methylophilaceae was the only family significantly positively associated with biomass. Members of this family have been associated with improved nutrient acquisition (Kumar et al., 2016), which may partly explain their positive association with plant biomass.

This study demonstrates that biowaste compost application has a significant influence on microbial community dynamics, enhancing diversity, metabolic potential, and plant growth-promoting (PGP) traits in soil, the rhizosphere, and root compartments. These effects are more evident in the rhizosphere, with lesser effects observed in bulk soil and roots, where we assume a strong influence from the soil microbiota and the plant’s control, respectively. The rhizosphere environment is crucial for plant health (Bakker et al., 2018; Klein et al., 2022; Ling et al., 2022) and can be vulnerable to a progressive decline in biodiversity due to the crisis of soil biodiversity associated with climate change (Fierer et al., 2009; Jansson and Hofmockel, 2020; Ibáñez et al., 2023).

Future research should investigate the long-term effects of biowaste compost on microbial communities and plant health under field conditions, as well as the interactions between microbes derived from biowaste compost and native soil and plant microbiota. Understanding these complex interactions is crucial for developing sustainable fertilization strategies that maximize agricultural productivity while minimizing environmental impact. The potential use of biowaste compost as bioinoculant which provide a set of potential beneficial strains is in line with what is suggested by Kaminsky and colleagues (Kaminsky et al., 2019), who suggest the use of consortia of microbes instead of single strains to overcome the difficulties of production, application, and stability of the inoculum in the soil.

By enriching soil and rhizosphere microbial communities with beneficial taxa and functional traits, biowaste compost has the potential to enhance soil health and to support plant growth and resilience. The increased abundance of plant growth-promoting taxa and metabolic pathways in biowaste compost-treated samples underscores its role in promoting sustainable agricultural practices.

## LIST OF ABBREVIATIONS

ACC deaminase: 1-aminocyclopropane-1-carboxylate deaminase
ASV: Amplicon sequence variant
BS: Bulk soil
BS.M42: Bulk soil+biowaste compost
BS.M42ST: Bulk soil+ sterile biowaste compost
BS.U: Bulk soil+Urea
C.M42: Batch number of biowaste compost used in the experiments
NMDS: Non-metric Multidimensional Scaling analysis
PGP: plant growth promoting
PCoA: Principal Coordinate Analysis
R.M42: Root+biowaste compost
R.M42ST: Root+sterile biowaste compost
R.U: Root+Urea
RH.M42: Rhizosphere+biowaste compost
RH.M42ST: Rhizosphere+sterile biowaste compost
RH.U: rhizosphere+Urea
SOIL: Soil

## Authors’ contributions

RC, GG, and NdB conducted experiments and analyzed the data. ML designed and optimized the functional analysis. SN conceptualized the research, designed the experimental plan, analyzed data, and wrote the manuscript. TB and WZ provided material, performed soil and biowaste compost chemical analysis, and funded the project. BB reviewed and edited the manuscript, supervised the project, and contributed to funding acquisition. All authors read and approved the final manuscript.

## Data availability statement

The sequence dataset generated and/or analysed during the current study is available in the Embl database under Project S-BSST2068, https://www.ebi.ac.uk/biostudies/studies/S-BSST2068?key=eeb5562a-77c5-46d8-a7f4-4bccbd7767ef). All the datasets generated by the analysis are available as supplementary material.

Supplementary Tables with original datasets are available on Zenodo platform (10.5281/zenodo.23717896).

## Acknowledgements

We thank Dr. Alberto Orgiazzi for assistance with functional data analysis presented in this paper, and Prof. Carlo Nicoletto for support in the selection of chemical fertilizer and its dose optimization. We thank the Erasmus Trainee Kalamara Vasileia from Democritus University of Thrace (Greece) and the BS student (BS in Natural Sciences, Dept. of Biology, University of Padova, Italy) Clarissa Laura Angiolin for the lab work.

## Additional files legends

**Fig S1**

PCoA and NMDS data visualization

PCoA analysis of the community of the sample groups and NMDS visualization per soil compartment. **(A)** Bray-Curtis and **(B)** Jaccard PCoA. BS.U2 is a technical replicate of BS.U used as a control for reproducibility. **(C)** NMDS and **(D)** PCoA of Jaccard distance matrix.

**Fig S2**

Rarefaction analysis for the alpha diversity indexes used in the community analysis.

**(A)**: rarefaction curve for Shannon index; **(B)** rarefaction curve of Pielou_e index; **(C)** rarefaction curve of Simpson index; **(D)** rarefaction curve for Dominant_features index, **(E)** rarefaction curve for Dominance index. The number of reads randomly chosen for normalization had a cutoff of 39787.

**Fig S3**

Barplot of the relative abundance for the first 10 families per single sample.

Complete overview per single sample of the relative abundance of the 10 most representative families.

**Fig S4**

Statistical comparison of the family abundance of BS.M42 vs BS.U and RH.M42 and RH.U A t-test is conducted to identify species with significant variation between groups (p-value < 0.05) at the family rank. In the left panel, the abundance of species showing significant differences between groups is displayed. Each bar represents the mean abundance for each group of species showing significant differences. The right panel presents the confidence interval for between-group variations. The leftmost part of each circle indicates the lower limit of the 95% confidence interval, while the rightmost part indicates the upper limit. The center of the circle represents the difference in the mean value. The color of the circle corresponds to the group with the higher mean value. The rightmost value indicates the p-value of the significance test for between-group variation. **(A)** comparison between BS.M42 vs BS.U and **(B)** comparison between RH.M42 and RH.U

**Fig S5**

Statistical comparison of family abundance between R.M42 and R.U and genus abundance between RH.M42 and RH.U A T-test is performed to determine species with significant variation between groups (p-value < 0.05) at the family **(A)** and genus **(B)** rank. In the left panel, there is the abundance of species showing significant differences between groups. Each bar represents the mean abundance of the species exhibiting these significant differences. The right panel shows the confidence interval for between-group variations. The leftmost part of each circle indicates the lower limit of the 95% confidence interval, while the rightmost part represents the upper limit. The center of the circle indicates the difference in mean value. The color of the circle correlates with the group whose mean value is higher. The rightmost value represents the p-value of the significance test for between-group variation. **(A)**: comparison of family abundance between R.M42 vs R.U; **(B)**: comparison of genus abundance between RH.M42 and RH.U.

**Fig S6**

Barplot of the relative abundance of Firmicutes genera in the sample groups. Complete overview per group of samples of the relative abundance of the 20 most representative genera within the Firmicutes phylum.

**Fig S7**

Statistical comparison of genus abundance between RH.M42ST and RH.U A T-test is performed to determine species with significant variation between groups (p-value < 0.05) at genus rank. In the left panel, there is the abundance of species showing significant differences between groups. Each bar represents the mean abundance of the species exhibiting these significant differences. The right panel shows the confidence interval for between-group variations. The leftmost part of each circle indicates the lower limit of the 95% confidence interval, while the rightmost part represents the upper limit. The center of the circle indicates the difference in mean value. The color of the circle correlates with the group whose mean value is higher. The rightmost value represents the p-value of the significance test for between-group variation.

**Fig S8**

Leaf dry biomass and its association with bacterial families in the tomato rhizosphere and roots. **(A)** Leaf dry biomass of plants treated with compost, sterile compost, or urea (n = 10). Different letters indicate significant differences according to pairwise Wilcoxon rank-sum tests with Benjamini-Hochberg correction. **(B–C)** Spearman correlations between leaf dry biomass and the relative abundance of the 30 most abundant bacterial families, selected independently in the rhizosphere (**B**) and root (**C**) compartments. Correlations were calculated separately for each treatment. Red indicates positive correlations, blue indicates negative correlations, and colour intensity represents the magnitude of Spearman’s ρ.

## Supplementary Tables

**Table S1**

Soil chemical/physical characteristics.

The table shows the chemical/physical characteristics of the agricultural soil used in this study. Methods of analysis are reported in the table and follow UNI/EU standards.

**Table S2**

Compost chemical/physical characteristics. The table shows the chemical/physical characteristics of the compost used in this study. Methods of analysis are reported in the table and follow UNI/EU standards.

**Table S3**

ADONIS analysis on the Bray-Curtis distance matrix

**Table S4**

ADONIS analysis on the Jaccard distance matrix

Table S5 Table reporting values of the diversity indexes Observed Features, Simpson and Shannon statistical analysis.

**Table S6**

Table reporting values of the evenness indexes Pielou and Dominance statistical analysis.

**Table S7**

Relative abundance of the 10 most representative phyla in the sample groups

**Table S8**

Relative abundance of the 10 most representative families in the samples

**Table S9**

T-test analysis results for the comparison BS.M42 vs BS.U

**Table S10**

T-test analysis results for the comparison RH.M42 vs RH.U

**Table S11**

T-test analysis results for the comparison R.M42 vs R.U

**Table S12**

T-test analysis results for the comparison RH.M42ST vs RH.U

**Table S13**

FEAST Composition matrix and group differences tested using the Wilcoxon rank-sum test

**Table S14**

T-test analysis results for the comparison of predicted pathways on BS.M42 vs BS.U

**Table S15**

T-test analysis results for the comparison of predicted pathways on R.M42 vs R.U

**Table S16**

T-test analysis results for the comparison of predicted pathways on RH.M42 vs RH.U

**Table S17**

Pathway annotation for all the sample groups

**Table S18**

T-test analysis results for the comparison of predicted enzymatic activities on RH.M42 vs RH.U

**Table S19**

Spearman correlations between leaf dry biomass and bacterial family relative abundance in tomato rhizosphere and root compartments

