## Supplementary figures and images for "Compost from urban biowaste shapes the bacterial community and its functional traits in the rhizosphere of tomato plants"

### FIGS1

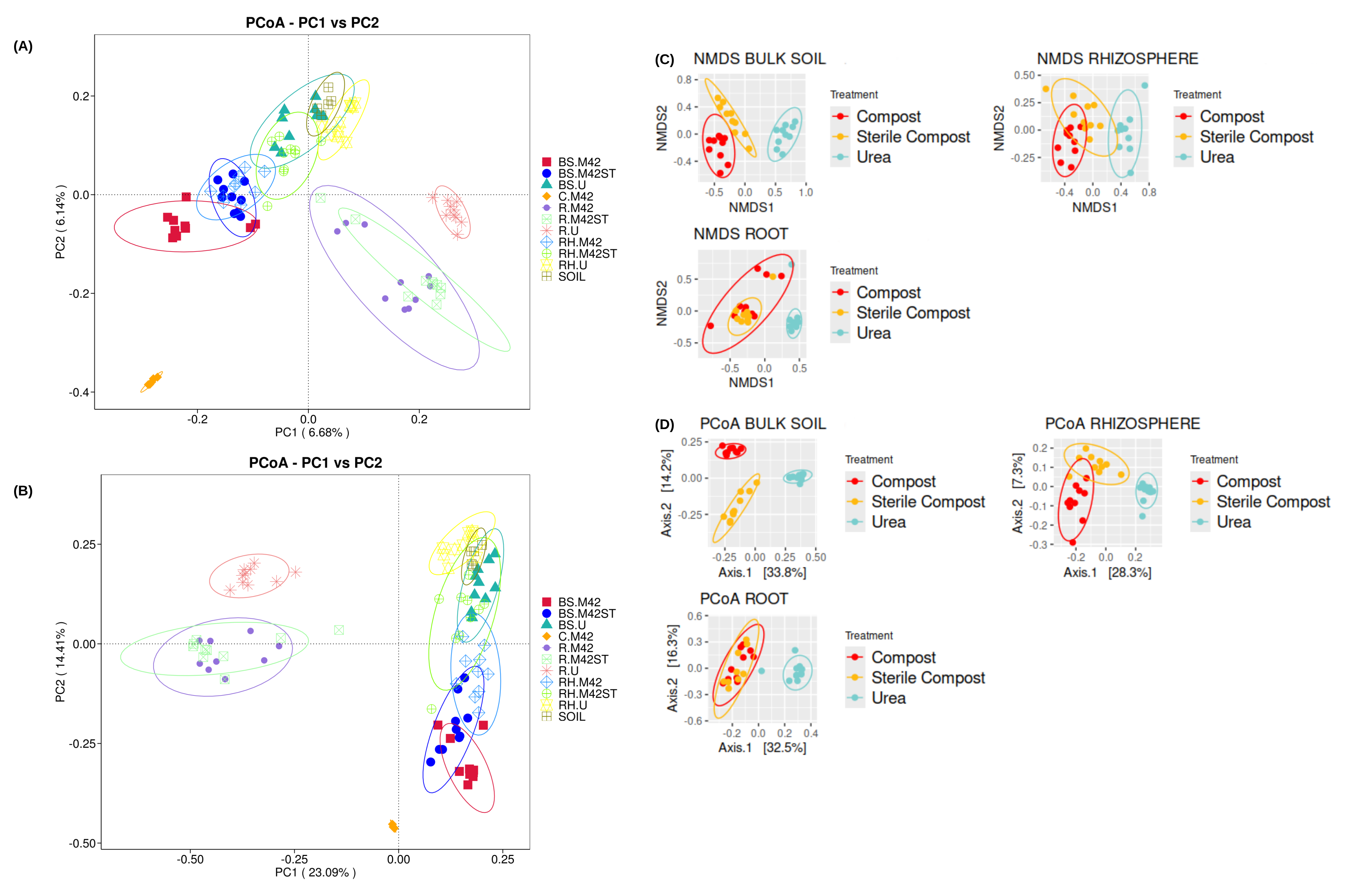

### FIGS2

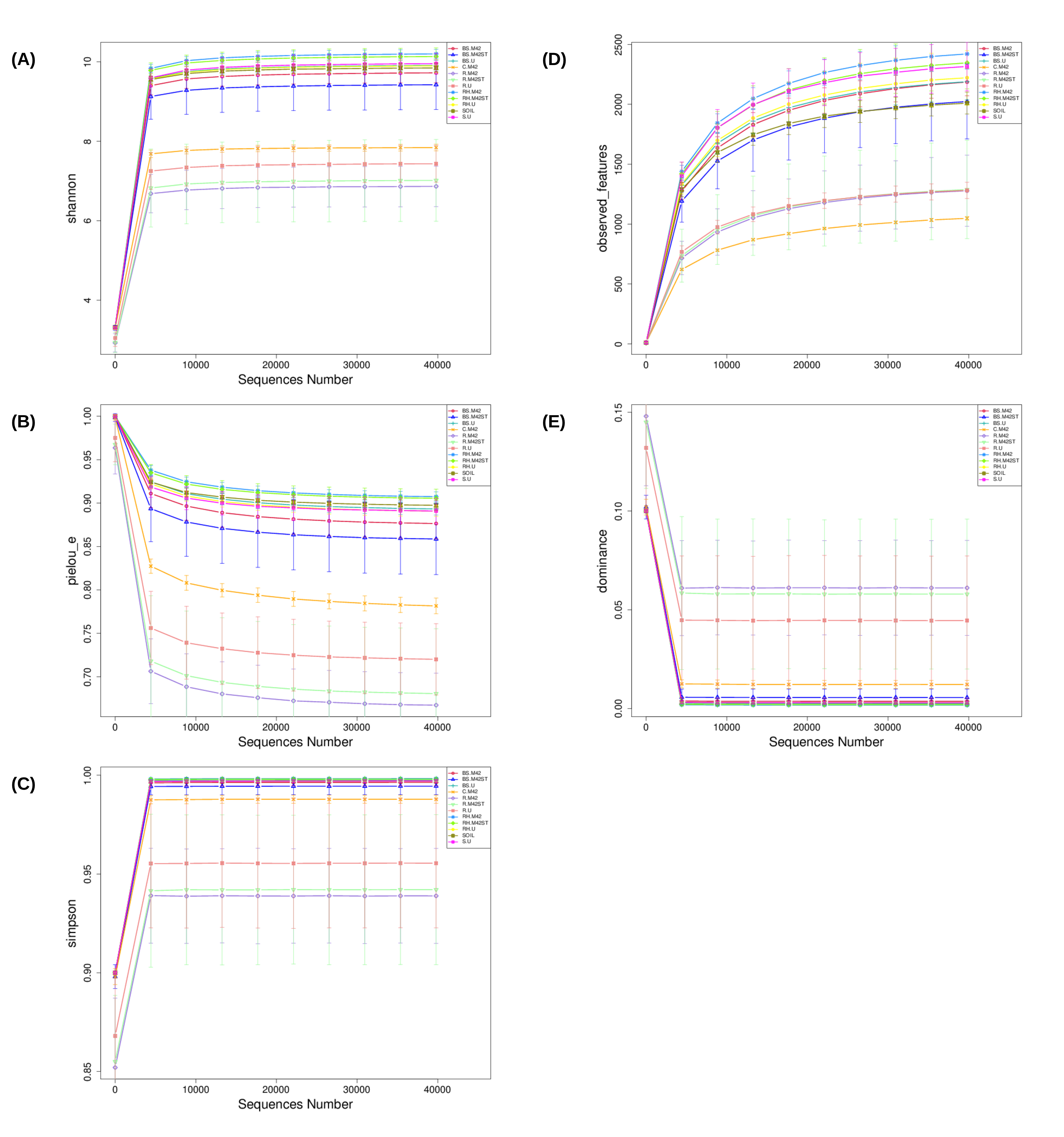

### FIGS3

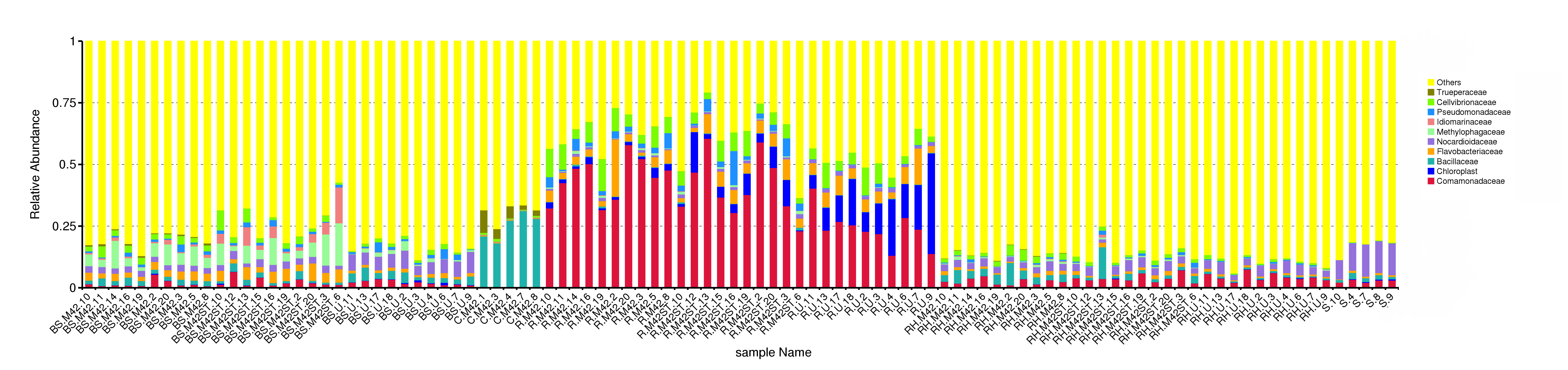

### FIGS4

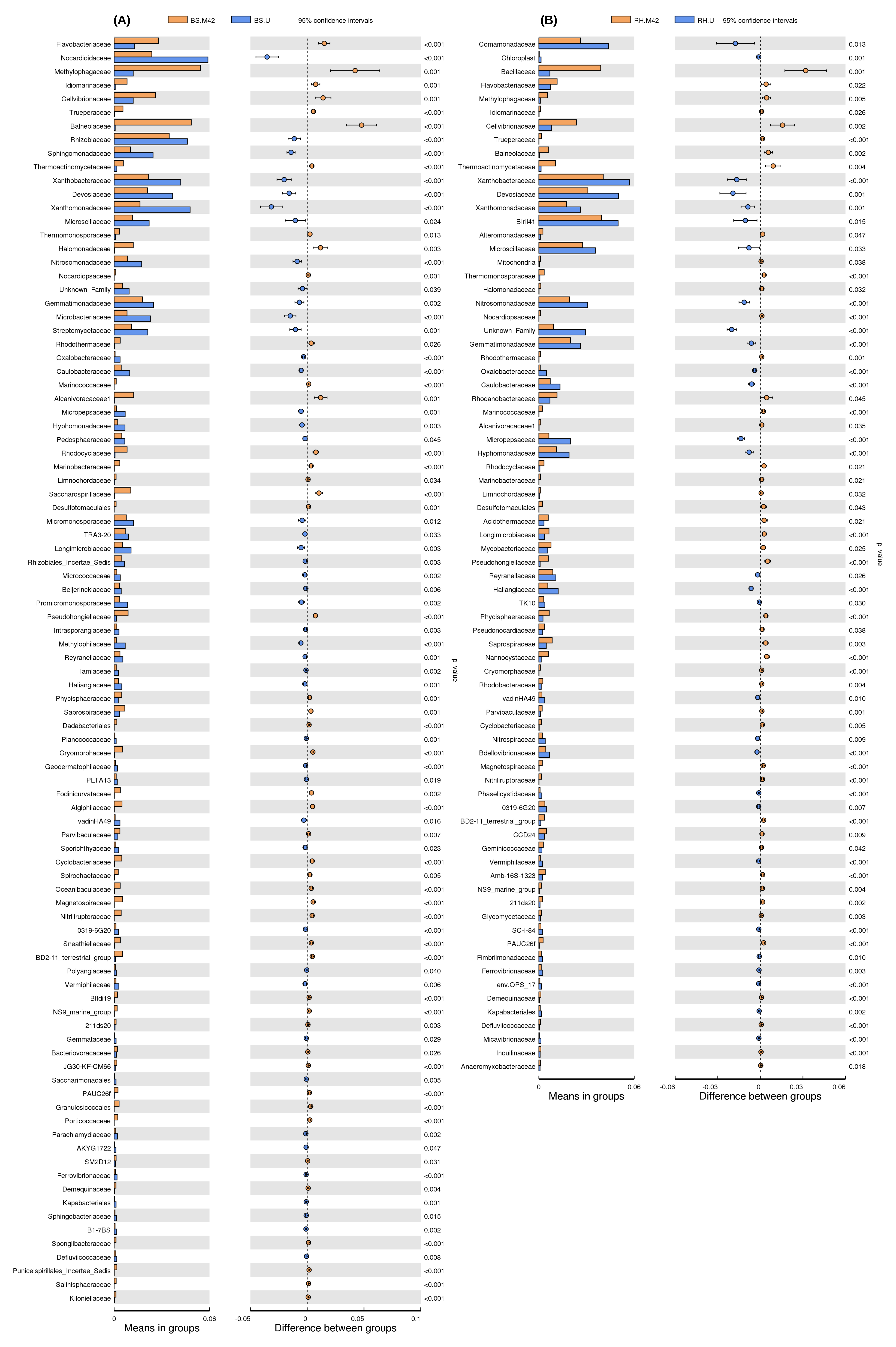

### FIGS5

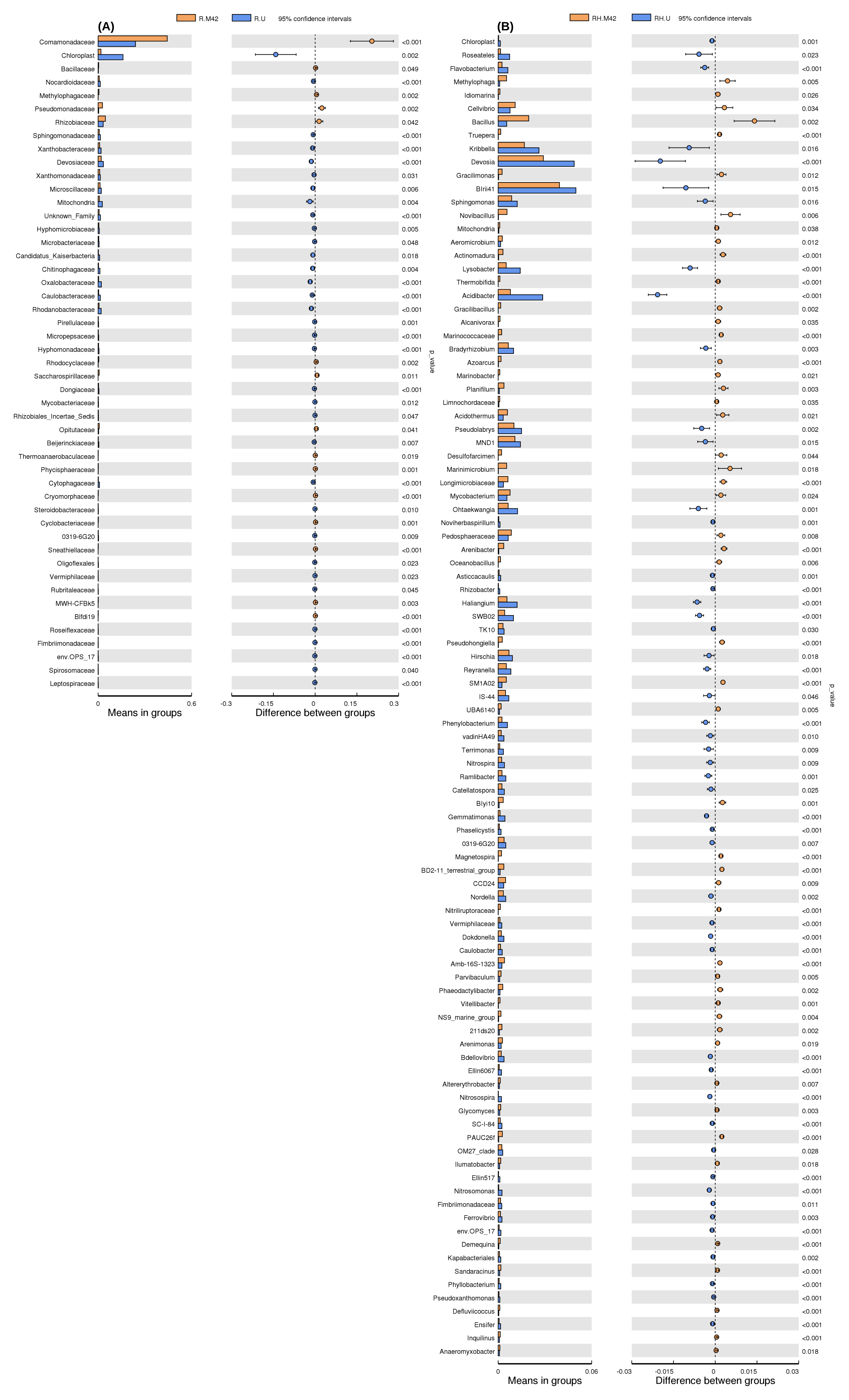

### FIGS6

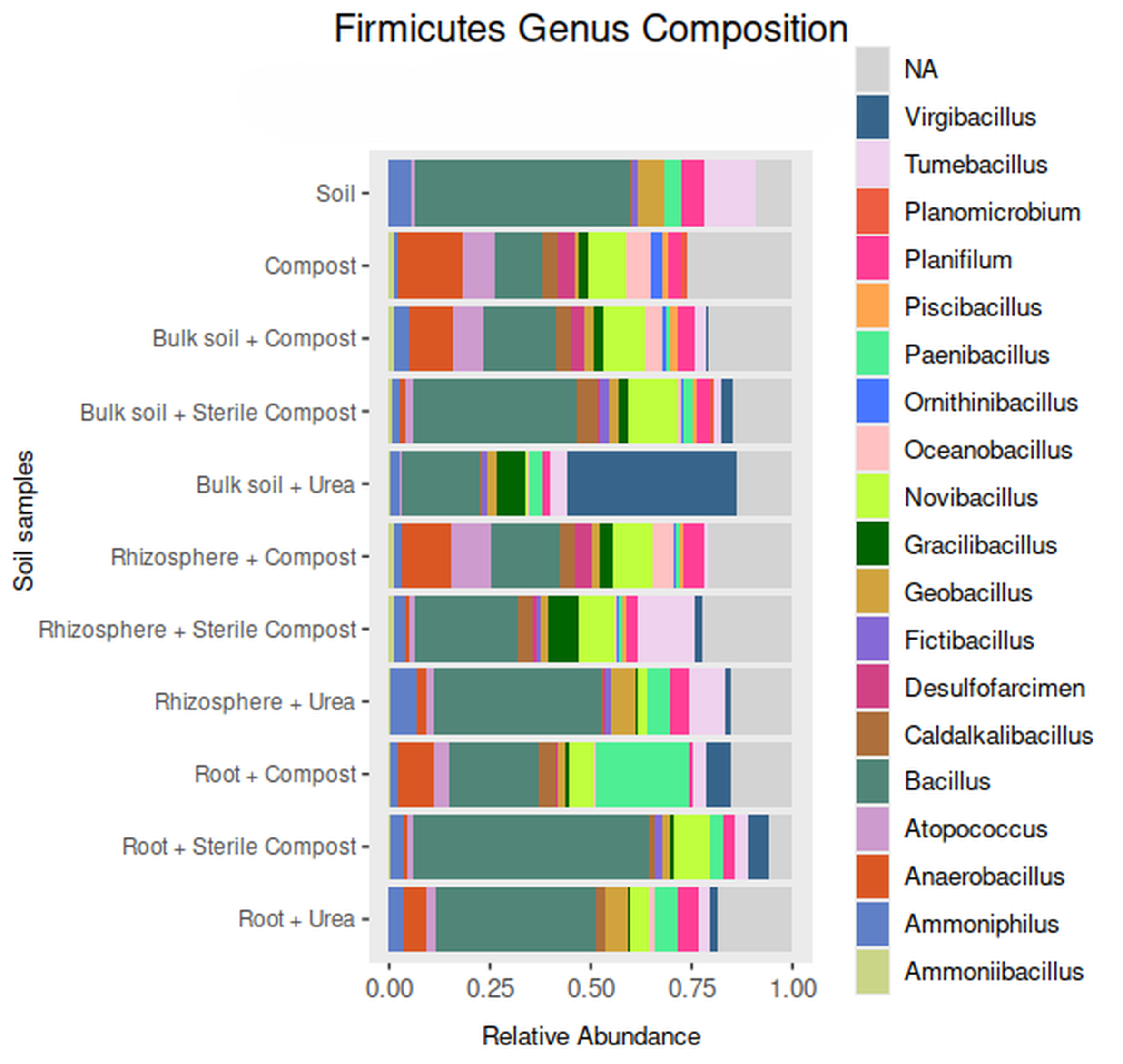

### FIGS7

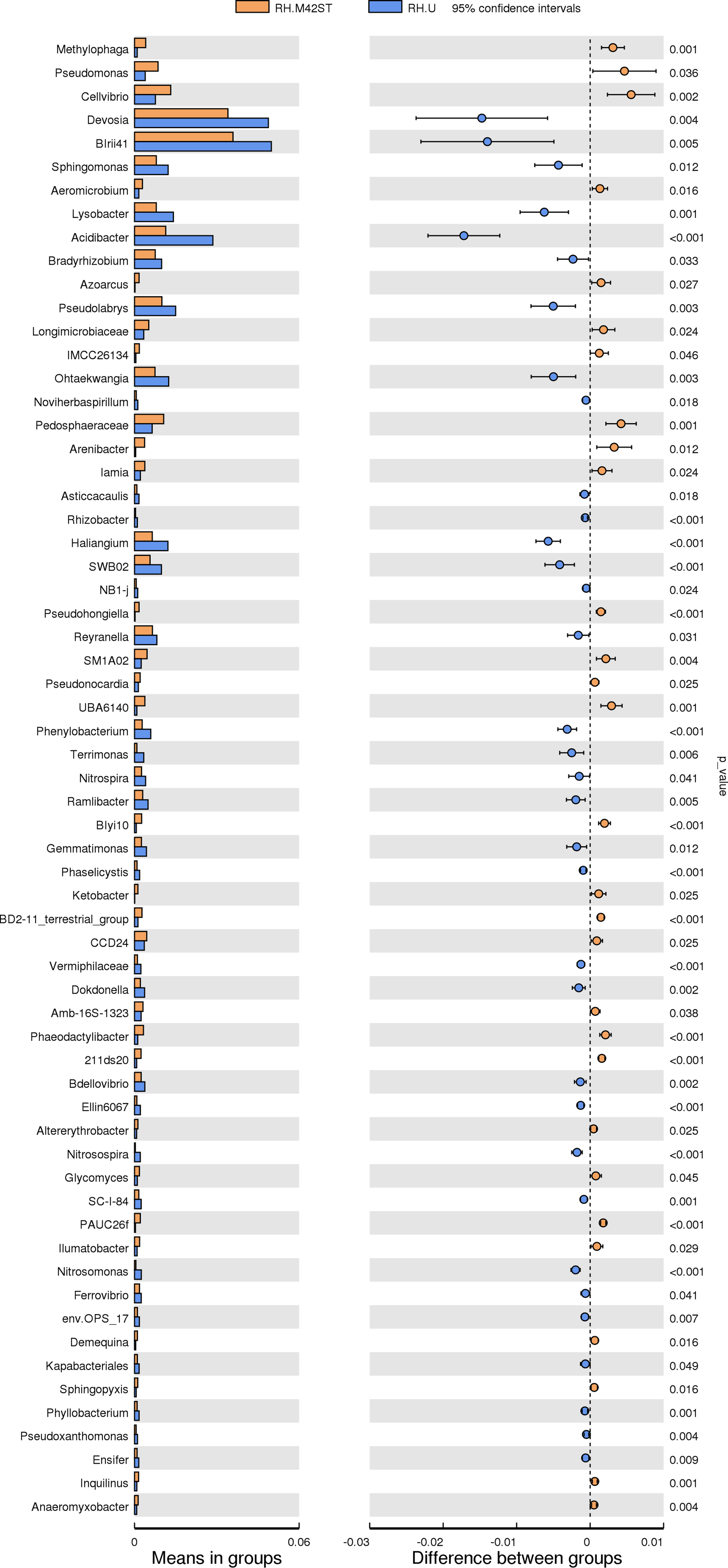

### FIGS8

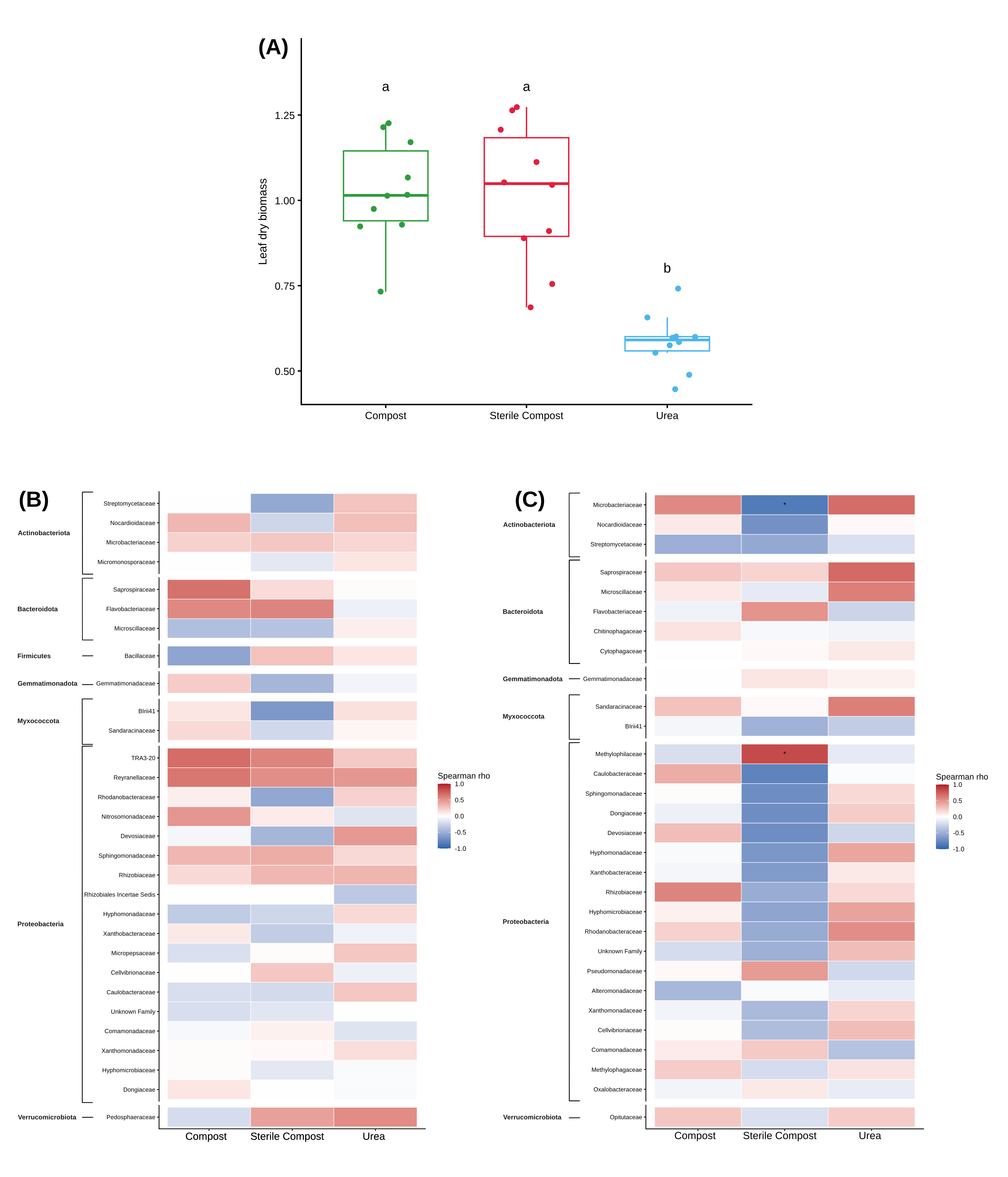
